# A previously missed population of antigen-specific CD8 T cells divides in the blood after vaccination

**DOI:** 10.1101/391664

**Authors:** Sonia Simonetti, Ambra Natalini, Antonella Folgori, Stefania Capone, Alfredo Nicosia, Angela Santoni, Francesca Di Rosa

## Abstract

Although clonal expansion is a hallmark of adaptive immunity, the location(s) where antigen-responding T cells enter cell cycle and complete it have been poorly explored. This lack of knowledge stems partially from the limited experimental approaches available. By using Ki67 plus DNA staining and a novel data analysis technique, we distinguished antigen-specific CD8 T cells in G_0_, in G_1_, and in S-G_2_-M phases after intramuscular vaccination of BALB/c mice with antigen-expressing viral vectors. We discovered an entire population of cycling cells that are usually missed. This “extra” population was present early after vaccination in lymph nodes, spleen and, surprisingly, also in the blood, which is not expected to be a site for mitosis of normal non-leukemic cells. These results have implications for previous and future immunological studies in animal models, and potentially in humans. They might also inspire hematologists to seek for other missed populations of dividing cells in blood.

## Introduction

Cell-to-cell interactions within tissue niches in solid organs and hematopoietic bone marrow (BM) regulate proliferation of stem cells and differentiated progenitors (1, 2), along with structural, physical, paracrine and neural cues provided by the microenvironment (3). Similarly, clonal expansion of T cells during adaptive immune responses is driven by antigen presenting cells within specialized niches in lymphoid organs, where local chemokines and cytokines guide T cell responses (4).

We nevertheless still lack essential spatial information on clonal expansion, particularly as to the location of T cells during each phase of the cell cycle. To date, T cell expansion in animal models has been mostly measured by dyes that label cells proliferating over time (e.g. CFSE; BrdU) (5, 6), without the ability to assess whether the labeled cells found in a particular location proliferated locally or rather migrated into that organ after dividing elsewhere. Another common method is staining for the intranuclear protein Ki67, after cell fixation and permeabilization (7–10). Though Ki67 is generally considered to label dividing cells, it actually labels all cells not in G_0_, not distinguishing actively cycling cells committed to mitosis (those in S-G_2_-M) from those in G_1_, which may quickly proceed into S, or stay in a prolonged G_1_, or even revert to G_0_ without dividing (11).

Here we used Ki67 plus DNA staining to track rare naïve antigen-specific CD8 T cells responding to vaccination in wild-type mice (12, 13). The naïve CD8 T cells clonally expanded, and we analyzed the resulting polyclonal population.

We found a significant number of antigen-responding CD8 T cells cycling in lymph nodes (LNs), spleen and (surprisingly) in the blood, a finding that opens new directions for the analysis of immune responses.

## Results

BALB/c mice were vaccinated intramuscularly (i.m.) against the model antigen, HIV-1 gag, using a recombinant chimpanzee-derived adenoviral vector (ChAd3-gag) and a Modified Virus Ankara (MVA-gag) for priming and boosting, respectively (12). The cell cycle stages of gag-specific CD8 T cells were analyzed using Hoechst 33342, a DNA dye, and anti-Ki67 mAb (14, 15).

Fig 1A-B shows the steps for analysing gag-specific CD8 T cells by flow cytometry, fig. 1B an example of spleen and LN cell analysis at day (d) 3 post-boost. Steps 1-2 identify single cells by DNA analysis, and live cells by dead cell marker exclusion. Step 3 uses Forward Scatter-A (FSC-A) and Side Scatter-A (SSC-A) profiles to identify certain leukocyte populations. Lymphocytes tend to have low SSC-A and medium-low FSC-A, whereas granulocytes have high SSC-A, and are normally excluded from the canonical ‘narrow’ gate used for lymphocyte studies (16–19) (Fig. 1B, Step 3, ‘narrow’). However, we noticed an unusual population of cells with high SSC-A that appeared only in vaccinated spleens and contained a significant number of antigen-specific lymphocytes (Fig. 1B, Step 3, arrow). When we enlarged our FSC-A/ SSC-A gate (Step 3 ‘relaxed’), before labeling CD8 T cells (Step 4) and antigen-specific T cells (Step 5), we found a 2-6 fold greater proportion of gag-specific CD8 T cells in the ‘relaxed’ gate population than in the ‘narrow’ gate population: not only in the spleen but also in the LNs (Fig. 1B-D). Although this gating strategy is novel for standard ex vivo studies of lymphocytes (Fig. 1-S1), cells with high FSC-A and high SSC-A are often included when examining in vitro activated T cells (20).

**FIGURE 1.**
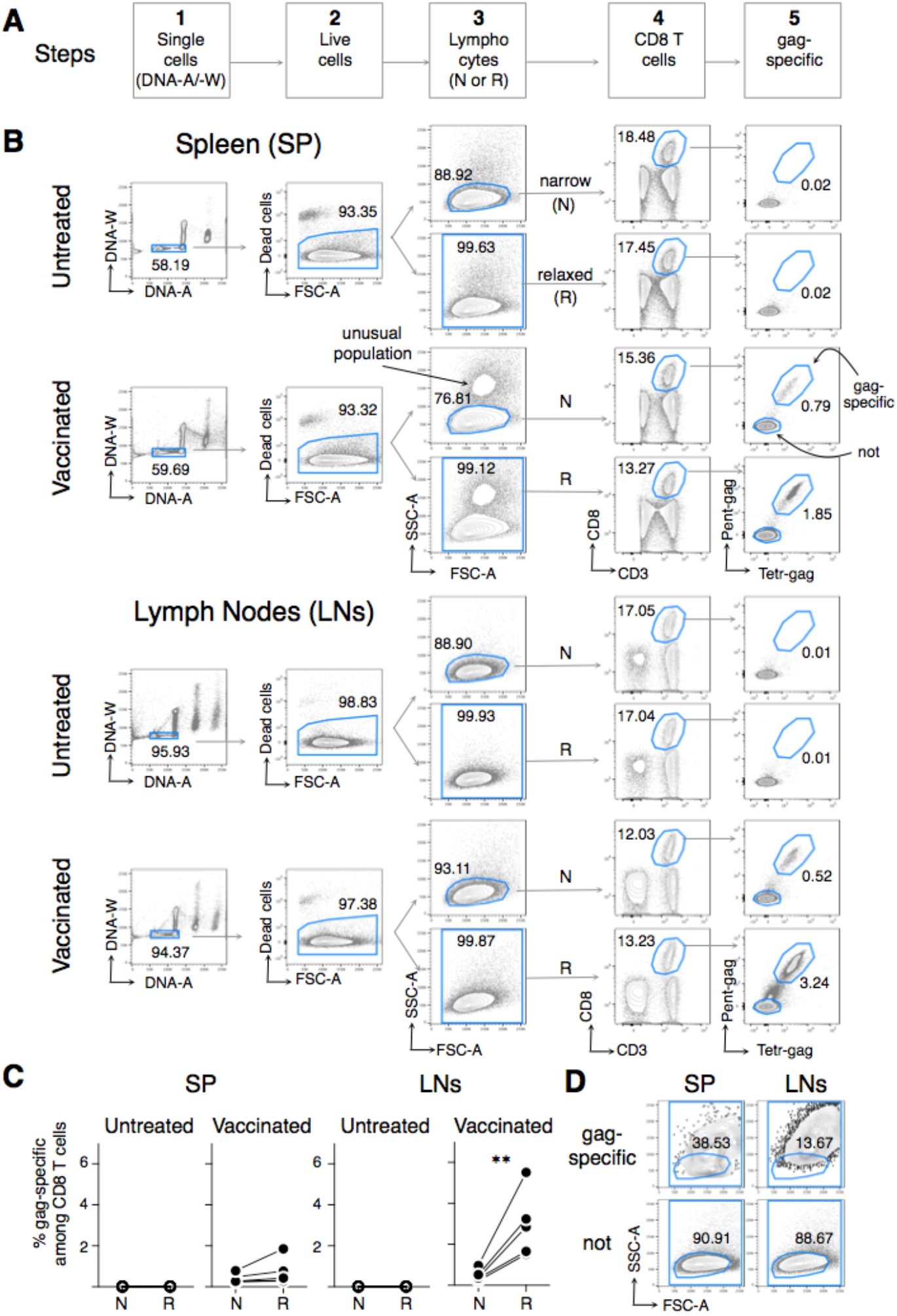
Comparison between the narrow (N) and the relaxed (R) gating strategy to evaluate frequency of gag-specific CD8 T cells from spleen and LNs of vaccinated mice at day (d) 3 post-boost. Female BALB/c mice were vaccinated by prime i.m. with ChAd3-gag (10^7^ vp) and boost i.m. with MVA-gag (10^6^ pfu). Cells from Spleen (SP) and LN (LNs) of vaccinated and untreated mice were analyzed by flow cytometry at d3 post-boost. CD3^(−)^ cells were gated out when acquiring spleen samples. (A) Scheme of the gating strategy for analysis of flow cytometry data in 5 steps, to identify the following cells: single cells (Step 1); live cells (Step 2); lymphocytes (Step 3); CD8 T cells (Step 4); gag-specific cells (Step 5). (B) Examples of flow cytometry analysis of cells from spleen (top) and LNs (bottom). At step 1, we discriminated single cells from doublets and aggregates by DNA content (DNA-A versus DNA-W). At Step 2 we excluded dead cells by using the eFluor780 Fixable Viability Dye. At Step 3, we used either the canonical gate for lymphocyte analysis (‘narrow’, N) or our proposed gate (‘relaxed’, R) in the FSC-A/ SSC-A plot, as indicated. At Step 4 we gated on CD3^+^ CD8^+^ cells, and at Step 5 we evaluated the percentages of gag_197-205_ (gag)-specific cells among them, by combined staining with Pent-gag and Tetr-gag. The numbers represent the percentages of cells in the indicated regions. The arrow in the vaccinated spleen FSC-A/ SSC-A plot indicates an unusual population of cells that was excluded by the N gate (see main text). (C) Summary of gag-specific CD8 T cell frequencies in spleen and LNs. The figure summarizes results obtained in 5 prime/boost experiments with a total of 30 mice. Statistically significant differences between N and R are indicated (** *p* ≤ 0.01). Differences in the frequency of gag-specific CD8 T cells between untreated and vaccinated mice were statistically significant both in spleen and LNs, using either R or N gating strategy (*p* ≤ 0.05, not shown). (D) Typical FSC-A/ SSC-A plots of gag-specific and not-specific CD8 T cells from spleen and LNs of vaccinated mice at d3 post-boost, gated using the R gate as in B.

In order to discriminate between gag-specific CD8 T cells in G_0_, G_1_, and S-G_2_-M, we examined Ki67 expression plus DNA content, using either the ‘narrow’ or the ‘relaxed’ gate (Fig. 2). We observed a striking difference in the percentages of proliferating cells between the two strategies. The ‘narrow’ gate missed most of the dividing cells in S-G_2_-M (<2%), whereas the ‘relaxed’ gate revealed that these cells made up to 42% of the gag-specific cells in LNs and 26% in spleen (Fig. 2A, C). Cell cycle entry and progression was accompanied by a graded increase of FSC-A, and more prominently of SSC-A (Fig. 2B-C). Proliferation was also seen after a single priming dose, though the kinetics were slower and there were fewer cells in S-G**2**-M (Fig. 2-S1).

**FIGURE 2.**
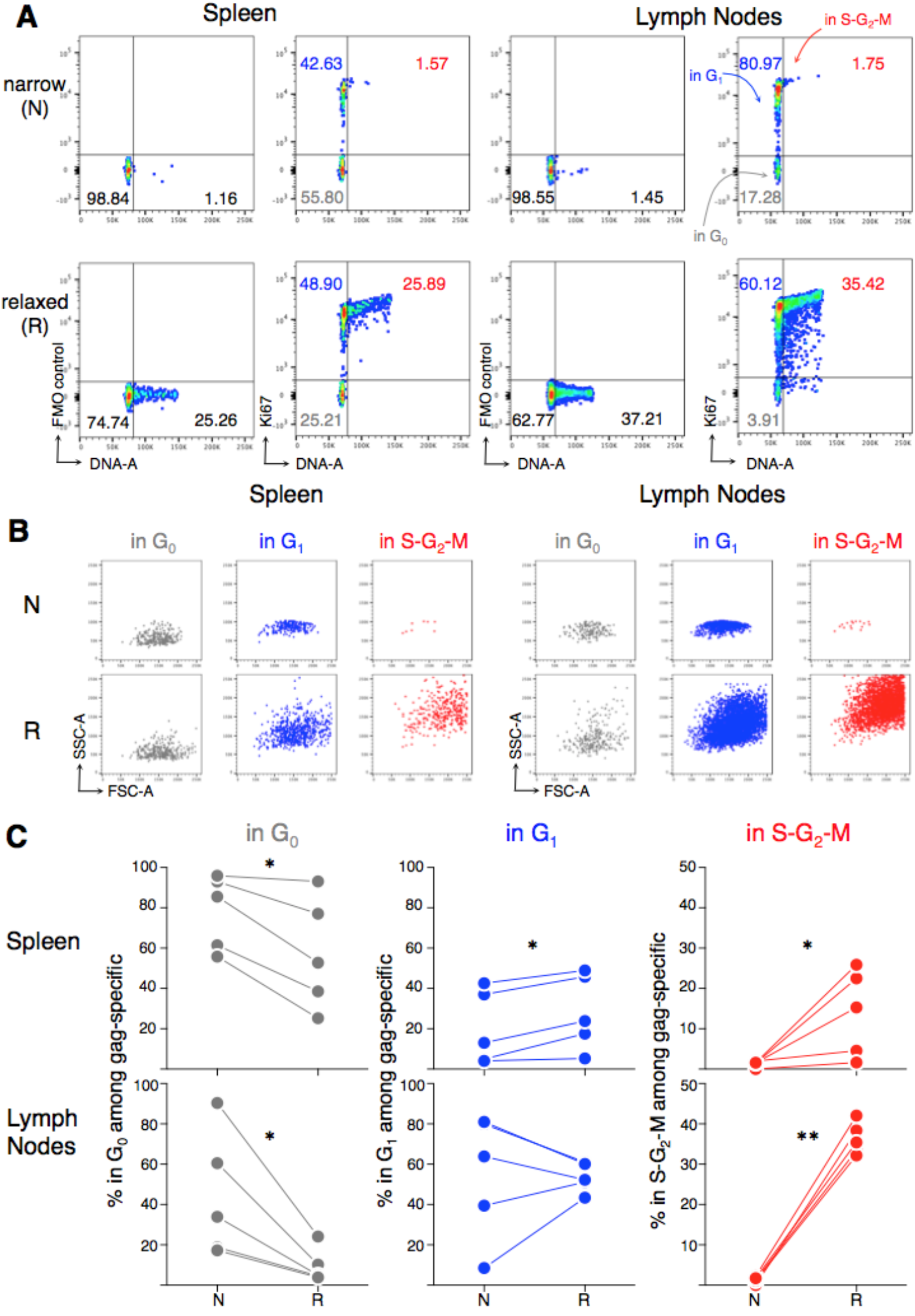
Comparison between the narrow (N) and the relaxed (R) gating strategy to evaluate cell cycle of gag-specific CD8 T cell from spleen and LNs of vaccinated mice at d3 post-boost. Cell cycle of gag-specific CD8 T cells at d3 post-boost was analyzed by Ki67 plus DNA staining, using either the N or the R gating strategy as in Fig. 1B. **(A)** Typical DNA/ Ki67 staining profiles of spleen (left) and LNs (right), after gating on gag-specific CD8 T cells. Fluorescence Minus One (FMO) controls and Ki67 staining are shown, as indicated. Based on DNA and Ki67 staining, cells in the following phases of cell cycle were identified in the corresponding quadrant: cells in G_0_ (Ki67^(−)^, 2n DNA), cells in G_1_ (Ki67^+^, 2n DNA) and cells in S-G_2_-M (Ki67+, 2n<DNA<4n), as indicated. The numbers represent the percentages of cells in the corresponding quadrant. **(B)** Typical FSC-A/ SSC-A plots of gag-specific CD8 T cells in G_0_, G_1_ and S-G_2_-M, gated as in A. **(C)** Summary of the percentages of gag-specific CD8 T cells in G_0_, in G_1_ and in S-G_2_-M in spleen (top) and LNs (bottom), gated as in A. The figure summarizes results obtained in 5 boost experiments with a total of 30 mice. Statistically significant differences between N and R are indicated (* *p* ≤ 0.05; ** *p* ≤ 0.01).

The ‘narrow’ gate missed up to a third of gag-specific CD8 T cells in the blood (Fig. 3A), which —with the ‘relaxed’ gate— averaged 2% at d3, 36% at d7, and 13% at d44 post-boost (Fig. 3B). As expected (12), gag-specific cells down-modulated CD62L (Fig 3-S1A-B). A well-defined population of mitotic gag-specific CD8 T cells was revealed uniquely using the ‘relaxed’ gate. Cells in S-G_2_-M were obvious at d3 (up to 13%) and less evident at d7 when Ki67+ peaked (up to 94%), suggesting that Ki67+ cells (non G_0_) persist in blood after mitotic cells disappear (Fig. 3C-D; 3-S1C-D). By day 44, almost all gag-specific cells were in G_0_ (Fig. 3C), suggesting that they had mostly switched to a resting memory state. We also saw mitotic antigen-specific CD8 T cells in blood after a single priming shot of vaccine (Fig. 3-S2).

**FIGURE 3.**
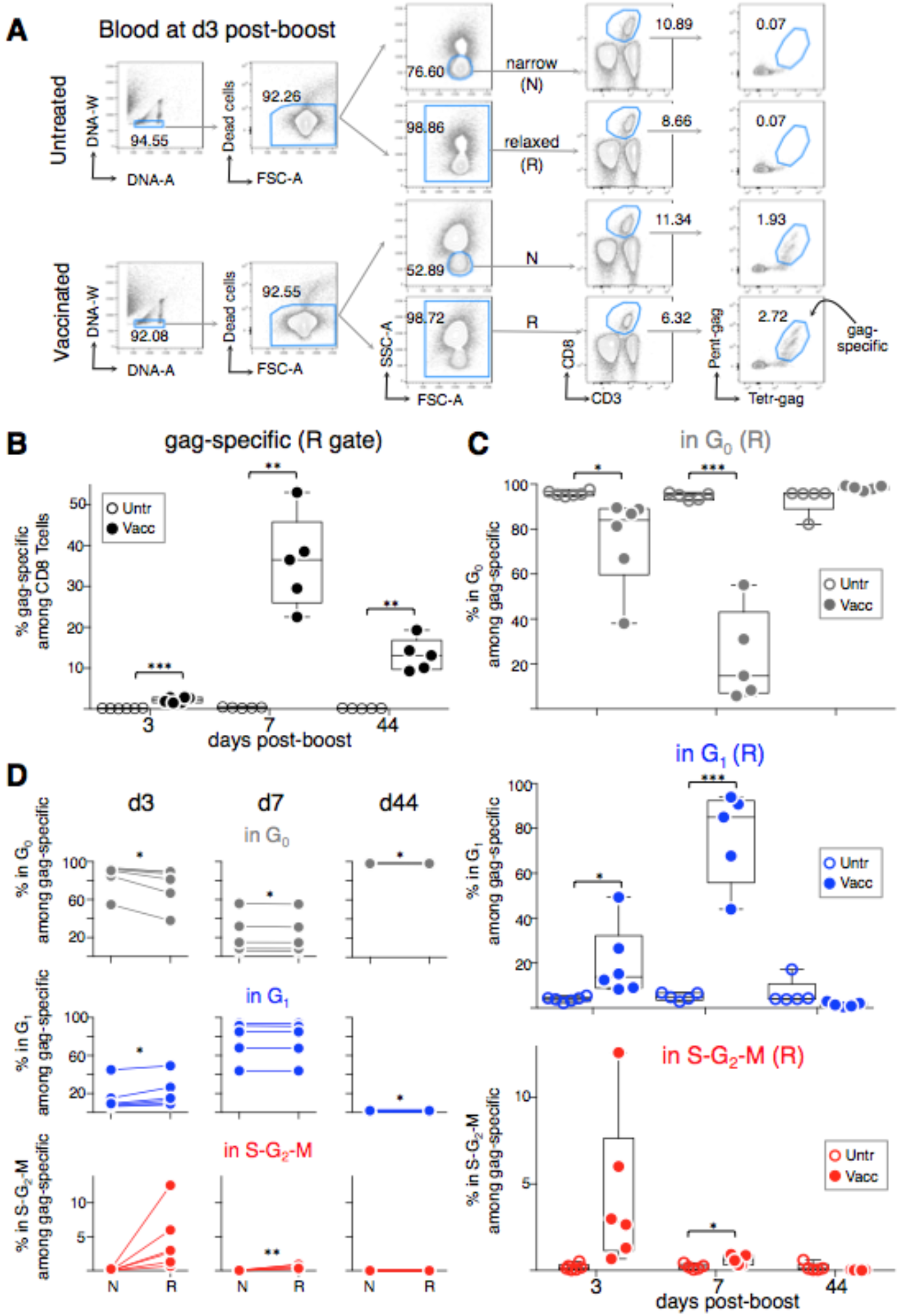
Analysis of the frequency and cell cycle of gag-specific CD8 T cells in the blood of vaccinated mice at d3, d7 and d44 post-boost. Female BALB/c mice were vaccinated as in Fig. 1. Blood was obtained from untreated and vaccinated mice at d3, d7 and d44 post-boost and gag-specific CD8 T cells were analyzed in 5 steps as in Fig. 1A and B, using either the N or the R gates at Step 3. **(A)** Example of flow cytometry analysis of blood cells from untreated and vaccinated mice at d3 post-boost. The numbers represent the percentages of cells in the indicated regions. **(B)** Summary of gag-specific CD8 T cell frequencies in the blood of untreated and vaccinated mice, obtained using the R gate. **(C)** Summary of the percentages of gag-specific CD8 T cells in G_0_ (top), in G_1_ (middle) and in S-G_2_-M (bottom) in the blood of vaccinated mice, compared with corresponding percentages among blood CD8 T cells from untreated controls, all obtained using the R gate. **(D)** Summary of the percentages of blood gag-specific CD8 T cells in G_0_ (top), in G_1_ (middle) and in S-G_2_-M (bottom) at d3, d7, and d44 post-boost, gated using either the N or the R gates as in A (see examples of cell cycle analysis using the R gate in Fig S3.1). The figure summarizes results obtained in 6 prime/boost experiments with a total of 60 mice. In B and C statistically significant differences between vaccinated and untreated mice are indicated at each time of analysis. In D statistically significant differences between N and R are indicated (\**p* ≤ 0.05; ** *p* ≤ 0.01; *** *p* ≤0.001).

Hypothesizing that the increased DNA content of the expanding CD8 T cells could be exploited as a marker to identify antigen-responding cells in the blood, we focussed on CD62L^(−)^ cells, as CD62L is generally down-regulated upon activation (Fig. 3-S1A-B). We evaluated the frequency of gag-specific cells among the following 4 populations of CD8 T cells (Fig. 4A): 1) total CD8 T (including naïve, memory and recently activated cells), 2) CD62L^(−)^ (non-naïve cells), 3) CD62L^(−)^ Ki67+ (non-G_o_ non-naïve cells), 4) CD62L^(−)^ in S-G_2_-M (dividing non-naïve cells). At day 3, the average percentage of gag-specific cells among the dividing non-naïve cells was 15-fold higher than among total CD8 T cells (Fig. 4C), sometimes up to 70% (Fig 4B), a much higher proportion than observed among the other 3 populations (Fig 4B-C). By d7, the gag-specific cells comprised 40% of the dividing non-naïve and 84% of the non-G_0_ non-naïve population. By d44 gag-specific cells were decreased in all the populations, though less evidently in the CD62L^(−)^ population (Fig. 4C). Results were similar, though the kinetics slower, after a single priming dose (Fig 4-S1).

**FIGURE 4.**
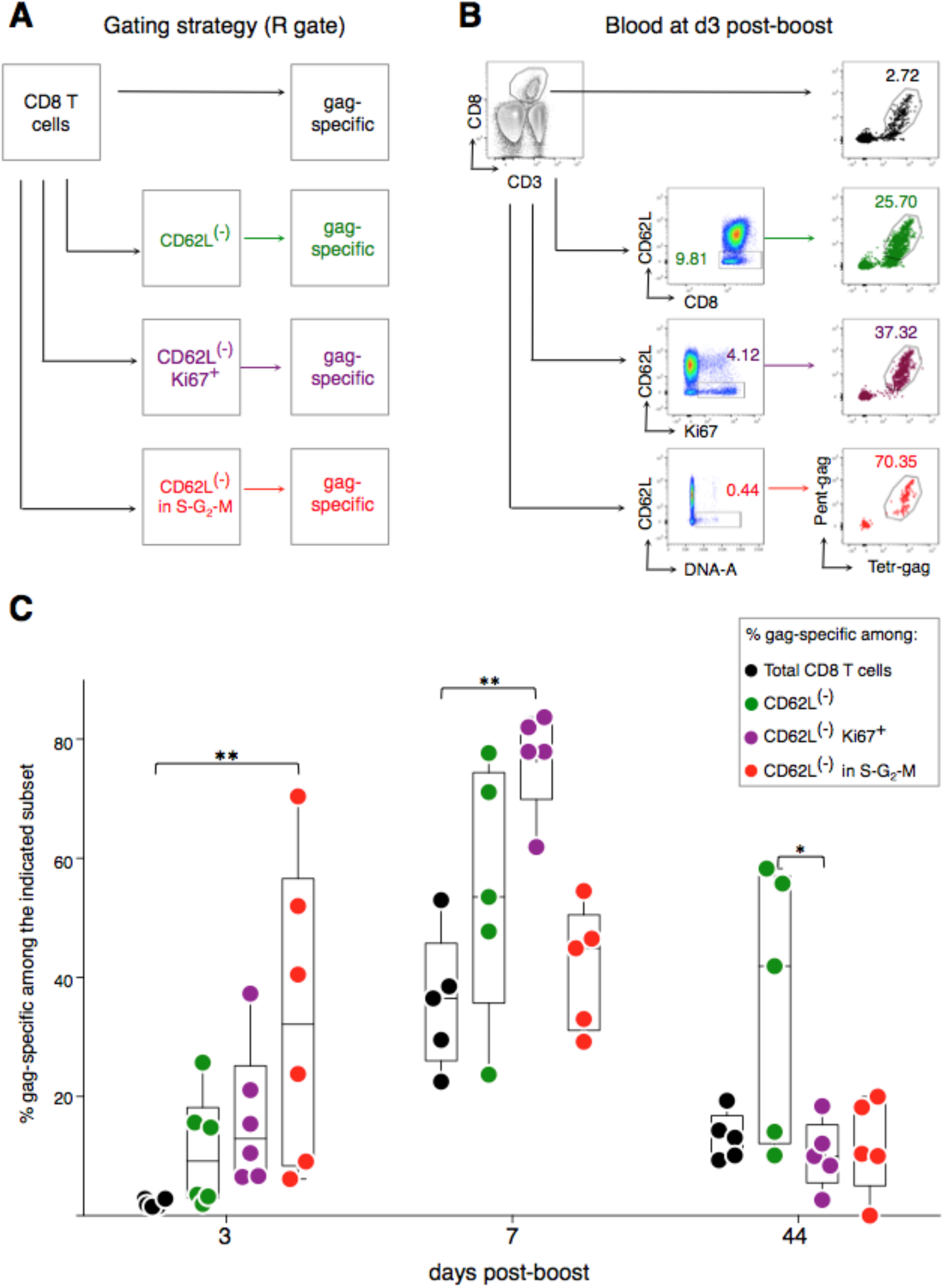
Specific enrichment of gag-specific CD8 T cells within a population of CD62L^(−)^ CD8 T cells in S-G_2_-M in the blood of vaccinated mice at d3 post-boost. Mice were vaccinated and blood samples analyzed at d3, d7 and d44 post-boost as in Fig. 3, using the R gate. The frequency of gag specific CD8 T cells among the following cell populations was determined: total CD8 T cells; CD62L^(−)^ CD8 T cells; Ki67+ CD62L^(−)^ CD8 T cells, and CD62L^(−)^ CD8 T cells in S-G_2_-M. **(A)** Gating strategy. **(B)** Example of flow cytometry profiles at d3 post-boost. **(C)** Summary of the results. In B the numbers represent the percentages of cells in the indicated regions. In C statistically significant differences are indicated at each time of analysis (\**p* ≤ 0.05; ** *p* ≤ 0.01). The figure summarizes results obtained in 6 prime/boost experiments with a total of 60 mice.

Since CD62L is a cell membrane molecule, and DNA can be visualized using vital dyes, our results suggest that the dividing CD62L^(−)^ CD8 T cells in blood could potentially be a valuable source of live antigen-specific CD8 T cells at early times of response.

## Discussion

Long ago, Sprent showed that, within days of an immunogenic stimulus, antigen-specific ‘blast’ T cells circulated in the thoracic duct lymph (21). There was no way of knowing at that time whether these were proliferating or simply activated cells. Here we show that mitotic antigen-specific T cells circulate in the blood stream, challenging the current view that the S-G_2_-M phases of clonal expansion occur only in lymphoid organs, or sometimes in BM, or in extra-lymphoid follicles in tissues (22, 23). Thus proliferation is not always limited to supportive tissues sites, but cells that have been stimulated in one organ can expand while circulating to other sites.

Mitotic gag-specific CD8 T cells were found in the blood after a single dose of ChAd3-gag, although they were fewer than after a boost with MVA-gag, possibly due to slower kinetics, and/or differences in spatial distribution of antigen-responding CD8 T cells inside the LNs (24) that were reflected in the blood. Further studies will be necessary to elucidate whether naïve and memory cells behave differently upon stimulation in vivo, whether vaccination route matters, and whether the cycling CD8 T cell clones in the blood comprise a special highly dividing subset and/or express high affinity TCRs.

The majority of dividing CD8 T cells in blood, spleen and LNs showed increased FSC-A and unusually high SSC-A, likely due to changes in mitochondria, chromatin condensation, etc. (25, 26). Cells with these characteristics are usually excluded from the analysis of normal lymphocytes ex vivo, for example human blood lymphocytes in conditions apart from cancer. Considering that nearly all immunological studies in humans use blood samples, important informations are likely to be missed, perhaps leading to incorrect conclusions. For example, it was found that up to 70% of virus-specific CD8 T cells were Ki67+ in the blood of patients at early phases of primary infections (8, 9), whereas memory-phenotype CD8 T cells from the blood of donors with no apparent infections comprised about 2-10% of Ki67^+^ cells (27). Furthermore, an early increase of Ki67+ PD-1+ CD8 T cells was observed in the blood of a subset of lung cancer patients treated with checkpoint inhibitors, and it was proposed that this could be relevant for antitumor effects (28). In all these studies it was suggested that the Ki67^+^ cells were proliferating in response to a recent immunogenic stimulus (8, 9, 27, 28), however cells with high side scatter were discarded (9, 28), and DNA content was not evaluated (8, 9, 27, 28), thus it cannot be distinguished whether the Ki67^+^ were actively cycling, or rather they were non-proliferating cells in G_1_, possibly on their way back to G_0_. Furthermore, the proliferation —when present— was likely greatly underestimated. A single study in humans did use DNA staining, and found that an average of <0.1% of memory-phenotype CD8 T cells were in S-G_2_-M in the blood of donors with no systemic diseases (27). The interpretation at that time was that blood-derived memory CD8 T cells are resting (27, 29, 30). We suggest instead that the cells in S-G_2_-M could have been newly activated cells responding to an environmental antigen.

Our results have several potential translational uses. For example, human blood might be the source of enriched populations of recently activated CD8 T cells, proliferating in response to vaccines, infections, transplantation and cancers, that could be studied, cloned and used therapeutically, even without knowing the antigen to which they are responding, or as a way of searching for that antigen. And, in cases where a patient presents with symptoms of immune activation, but no obvious infection, an analysis of the mitotic cells in the blood could reveal clues as to the cause of the symptoms and/or the target of the response.

## Material and Methods

### Adenoviral and MVA vectors

Replication defective, ΔE1 ΔE2 ΔE3 ChAd3 vector encoding HIV-1 gag protein under HCMV promoter (ChAd3-gag, 21) (31) and Modified Vaccinia Ankara encoding the HIV-1 gag protein under the control of vaccinia p7.5 promoter (MVA-gag) were used in all experiments.

### Vaccination

Six-week-old female BALB/c mice from Envigo (S. Pietro al Natisone, Udine, Italy) were housed at Plaisant animal facility (Castel Romano, Rome, Italy), and divided into experimental groups of at least 40 mice each (untreated and vaccinated). All mice of the vaccinated group were primed with ChAd3-gag, and a subset was analyzed after priming only. The remaining primed mice were boosted once with MVA-gag, at either d60 (range 60-67) or d100 (range 95-109) post-prime. Results of d60 and d100 boosts were similar, thus we combined them. Viral vectors were administered i.m. in the quadriceps at a dose of 10^7^ viral particles (vp) for ChAd3-gag and 10^6^ plaque-forming units (pfu) for MVA-gag, in a volume of 50 μl per side (100 μl total). All experimental procedures were approved by the local animal ethics council and performed in accordance with national and international laws and policies (UE Directive 2010/63/UE; Italian Legislative Decree 26/2014). Vaccination procedures were performed under anesthesia, and all efforts were made to reduce animal numbers and minimize suffering.

### Organs

Spleen, LNs and blood were obtained at different times after either prime or boost, i.e. d7, d10 and d14 post-prime; d3, d7 and d44 post-boost. At each time, the organs were collected from 3 vaccinated and 3 untreated mice, and cells from the 3 mice of each group were pooled. At d7 (and in one experiment at d3) blood was obtained by submandibular vein puncture in conscious mice. At all the other time points organ harvesting was scheduled, thus blood was obtained by cardiac puncture upon carbon dioxide euthanasia. Blood was immediately put into Heparin or EDTA blood collection tubes and further processed for analysis. Single-cell suspensions were prepared from spleen and LNs (iliac and inguinal) by mechanical disruption and passage through cell strainers (32).

### Membrane Staining

Spleen and LN cells were incubated with Fixable Viability Dye conjugated with eFluor780 fluorochrome (Affimetrix, eBioscience, Santa Clara, CA) and background staining was blocked with anti-FcγR mAb (clone 2.4G2). Cells were then incubated for 15 minutes at 4°C with H-2k(d) AMQMLKETI APC-labeled Tetramer (Tetr-gag, NIH Tetramer Core Facility, Atlanta, GA) and PE-labeled Pentamer (Pent-gag, Proimmune, Oxford, UK) to stain for gag_197-205_(gag)-specific CD8 T cells. Cells were incubated for further 15 minutes at 4°C after addition of the following mAbs: anti-CD3 peridinin chlorophyll protein (PerCP)-Cy5.5 (clone 145-2C11, BD Biosciences), anti-CD8α BUV805 (clone 53-6.7, BD Biosciences), anti-CD62L phycoerythrin (PE)-Cy7 (clone MEL-14, Biolegend, San Diego, CA, USA). Blood samples were incubated for 30 minutes at RT with the above antibodies/reagents that were placed all together. After washing, blood cells were fixed with Cell Fix solution (BD Biosciences). Red cells were lysed with Pharm Lyse solution (BD Biosciences).

### Intracellular Staining

Intracellular staining for Ki67 and DNA was performed as previously described, with some modifications (14, 15). Cells were fixed and permeabilized with Foxp3/Transcription Factor Staining Buffer (Affimetrix, eBioscience). Intracellular staining was performed with anti-Ki67 mAb conjugated with Fluorescein isothiocyanate (FITC) or Alexafluor 700 (clone SolA-15; eBioscience). DNA was stained by incubation with Hoechst 33342 (Thermo Fisher Scientific, Waltham, MA).

### Flow cytometry analysis

Samples were analyzed by LSRFortessa flow cytometer (BD Biosciences), gating out CD3^(−)^ cells when acquiring spleen samples. Data were analysed using FlowJo software, v.10 (FlowJo, Ashland, OR, USA).

### Statistical analysis

At each time point, the vaccinated group was compared with its corresponding untreated group by performing a two-tailed unpaired Student t test with Welch’s correction. A two-tailed paired Student t test was used for comparison of N and R gates. Friedman test with Dunn’s multiple comparison was used for comparison of multiple cell subsets within vaccinated mice samples. Differences were considered significant when \**p* ≤ 0.05; ** *p* ≤ 0.01; *** *p* ≤0.001. Statistical analysis was performed using Prism v.6.0f, GraphPad Software (La Jolla, CA, USA).

### Author contributions

F.D. designed experiments, interpreted the results and wrote the paper with help by S.S., A. Natalini and A.S.; A.F., S. C. and A. Nicosia prepared the viral vectors and performed mouse treatments, S.S. and A.N. performed/analyzed flow cytometry experiments.

## Acknowledgements

We thank A. Hayday, M. Munoz-Ruiz and F. Granucci for discussion. A special thank to P. Matzinger for providing precious advices and for reading the manuscript. The following tetramer was obtained through the NIH Tetramer Facility: APC-conjugated H-2K(d) HIV gag 197–205 AMQMLKETI

[Figure 1-Figure Supplement 1: Comparison between the N and the R gating strategy to evaluate frequency of gag-specific CD8 T cells in spleen and LNs after single cell discrimination by FSC-A/ FSC-H].

[Figure 2-Figure Supplement 1. Analysis of frequency and cell cycle of gag-specific CD8 T cells in spleen and LNs after prime only].

[Figure 3-Figure Supplement 1. Examples of flow cytometry analysis of gag-specific CD8 T cells in the blood of vaccinated mice at d3, d7 and d44 post-boost.]

[Figure 3-Figure Supplement 2. Analysis of frequency and cell cycle of gag-specific CD8 T cells in the blood after prime only].

[Figure 4-Figure Supplement 1: Specific enrichment of gag-specific CD8 T cells within a population of CD62L^(−)^ CD8 T cells in S-G_2_-M in the blood of primed mice].

## References

1. Yamashita, Y. M. 2010. Cell adhesion in regulation of asymmetric stem cell division. Curr Opin Cell Biol 22: 605–610. doi: 10.1016/j.ceb.2010.07.009

2. Bianco, P. 2011. Bone and the hematopoietic niche: a tale of two stem cells. Blood 117: 5281–5288. doi: 10.1182/blood-2011-01-315069

3. Scadden, D. T. 2006. The stem-cell niche as an entity of action. Nature 441: 1075–1079. doi: 10.1038/nature04957

4. Castellino, F., A. Y. Huang, G. Altan-Bonnet, S. Stoll, C. Scheinecker, and R. N. Germain. 2006. Chemokines enhance immunity by guiding naive CD8+ T cells to sites of CD4+ T cell-dendritic cell interaction. Nature 440: 890–895. doi: 10.1038/nature04651

5. Murali-Krishna, K., J. D. Altman, M. Suresh, D. J. Sourdive, A. J. Zajac, J. D. Miller, J. Slansky, and R. Ahmed. 1998. Counting antigen-specific CD8 T cells: a reevaluation of bystander activation during viral infection. Immunity 8: 177–187. doi:

6. van Stipdonk, M. J., E. E. Lemmens, and S. P. Schoenberger. 2001. Naive CTLs require a single brief period of antigenic stimulation for clonal expansion and differentiation. Nat Immunol 2: 423–429. doi: 10.1038/87730

7. Pandrea, I., T. Gaufin, R. Gautam, J. Kristoff, D. Mandell, D. Montefiori, B. F. Keele, R. M. Ribeiro, R. S. Veazey, and C. Apetrei. 2011. Functional Cure of SIVagm Infection in Rhesus Macaques Results in Complete Recovery of CD4+ T Cells and Is Reverted by CD8+ Cell Depletion. PLoS Pathogens 7: e1002170. doi: 10.1371/journal.ppat.1002170

8. Zaunders, J. J. 2005. Early proliferation of CCR5+ CD38+++ antigen-specific CD4+ Th1 effector cells during primary HIV-1 infection. Blood 106: 1660–1667. doi: 10.1182/blood-2005-01-0206

9. van Aalderen, M. C., E. B. Remmerswaal, N. J. Verstegen, P. Hombrink, A. ten Brinke, H. Pircher, N. A. Kootstra, I. J. ten Berge, and R. A. van Lier. 2015. Infection history determines the differentiation state of human CD8+ T cells. J Virol 89: 5110–5123. doi: 10.1128/JVI.03478-14

10. Bolinger, B., S. Sims, L. Swadling, G. O’Hara, C. de Lara, D. Baban, N. Saghal, L. N. Lee, E. Marchi, M. Davis, E. Newell, S. Capone, A. Folgori, E. Barnes, and P. Klenerman. 2015. Adenoviral Vector Vaccination Induces a Conserved Program of CD8(+) T Cell Memory Differentiation in Mouse and Man. Cell Rep 13: 1578–1588. doi: 10.1016/j.celrep.2015.10.034

11. Di Rosa, F. 2016. Two Niches in the Bone Marrow: A Hypothesis on Life-long T Cell Memory. Trends Immunol 37: 503–512. doi: 10.1016/j.it.2016.05.004

12. Quinn, K. M., A. Da Costa, A. Yamamoto, D. Berry, R. W. Lindsay, P. A. Darrah, L. Wang, C. Cheng, W. P. Kong, J. G. Gall, A. Nicosia, A. Folgori, S. Colloca, R. Cortese, E. Gostick, D. A. Price, C. E. Gomez, M. Esteban, L. S. Wyatt, B. Moss, C. Morgan, M. Roederer, R. T. Bailer, G. J. Nabel, R. A. Koup, and R. A. Seder. 2013. Comparative analysis of the magnitude, quality, phenotype, and protective capacity of simian immunodeficiency virus gag-specific CD8+ T cells following human-, simian-, and chimpanzee-derived recombinant adenoviral vector immunization. J Immunol 190: 2720–2735. doi: 10.4049/jimmunol.1202861

13. Stanley, D. A., A. N. Honko, C. Asiedu, J. C. Trefry, A. W. Lau-Kilby, J. C. Johnson, L. Hensley, V. Ammendola, A. Abbate, F. Grazioli, K. E. Foulds, C. Cheng, L. Wang, M. M. Donaldson, S. Colloca, A. Folgori, M. Roederer, G. J. Nabel, J. Mascola, A. Nicosia, R. Cortese, R. A. Koup, and N. J. Sullivan. 2014. Chimpanzee adenovirus vaccine generates acute and durable protective immunity against ebolavirus challenge. Nat Med 20: 1126–1129. doi: 10.1038/nm.3702

14. Wilson, A., M. J. Murphy, T. Oskarsson, K. Kaloulis, M. D. Bettess, G. M. Oser, A. C. Pasche, C. Knabenhans, H. R. Macdonald, and A. Trumpp. 2004. c-Myc controls the balance between hematopoietic stem cell self-renewal and differentiation. Genes Dev 18: 2747–2763. doi: 10.1101/gad.313104

15. Hirche, C., T. Frenz, S. F. Haas, M. Döring, K. Borst, P. K. Tegtmeyer, I. Brizic, S. Jordan, K. Keyser, C. Chhatbar, E. Pronk, S. Lin, M. Messerle, S. Jonjic, C. S. Falk, A. Trumpp, M. A. G. Essers, and U. Kalinke. 2017. Systemic Virus Infections Differentially Modulate Cell Cycle State and Functionality of Long-Term Hematopoietic Stem Cells In Vivo. Cell Rep 19: 2345–2356. doi: 10.1016/j.celrep.2017.05.063

16. Cossarizza, A., H. D. Chang, A. Radbruch, M. Akdis, I. Andrä, F. Annunziato, P. Bacher, V. Barnaba, L. Battistini, W. M. Bauer, S. Baumgart, B. Becher, W. Beisker, C. Berek, A. Blanco, G. Borsellino, P. E. Boulais, R. R. Brinkman, M. Büscher, D. H. Busch, T. P. Bushnell, X. Cao, A. Cavani, P. K. Chattopadhyay, Q. Cheng, S. Chow, M. Clerici, A. Cooke, A. Cosma, L. Cosmi, A. Cumano, V. D. Dang, D. Davies, S. De Biasi, G. Del Zotto, S. Della Bella, P. Dellabona, G. Deniz, M. Dessing, A. Diefenbach, J. Di Santo, F. Dieli, A. Dolf, V. S. Donnenberg, T. Dörner, G. R. A. Ehrhardt, E. Endl, P. Engel, B. Engelhardt, C. Esser, B. Everts, A. Dreher, C. S. Falk, T. A. Fehniger, A. Filby, S. Fillatreau, M. Follo, I. Förster, J. Foster, G. A. Foulds, P. S. Frenette, D. Galbraith, N. Garbi, M. D. García-Godoy, J. Geginat, K. Ghoreschi, L. Gibellini, C. Goettlinger, C. S. Goodyear, A. Gori, J. Grogan, M. Gross, A. Grützkau, D. Grummitt, J. Hahn, Q. Hammer, A. E. Hauser, D. L. Haviland, D. Hedley, G. Herrera, M. Herrmann, F. Hiepe, T. Holland, P. Hombrink, J. P. Houston, B. F. Hoyer, B. Huang, C. A. Hunter, A. Iannone, H. M. Jäck, B. Jávega, S. Jonjic, K. Juelke, S. Jung, T. Kaiser, T. Kalina, B. Keller, S. Khan, D. Kienhöfer, T. Kroneis, D. Kunkel, C. Kurts, P. Kvistborg, J. Lannigan, O. Lantz, A. Larbi, S. LeibundGut-Landmann, M. D. Leipold, M. K. Levings, V. Litwin, Y. Liu, M. Lohoff, G. Lombardi, L. Lopez, A. Lovett-Racke, E. Lubberts, B. Ludewig, E. Lugli, H. T. Maecker, G. Martrus, G. Matarese, C. Maueröder, M. McGrath, I. McInnes, H. E. Mei, F. Melchers, S. Melzer, D. Mielenz, K. Mills, D. Mirrer, J. Mjösberg, J. Moore, B. Moran, A. Moretta, L. Moretta, T. R. Mosmann, S. Müller, W. Müller, C. Münz, G. Multhoff, L. E. Munoz, K. M. Murphy, T. Nakayama, M. Nasi, C. Neudörfl, J. Nolan, S. Nourshargh, J. E. O’Connor, W. Ouyang, A. Oxenius, R. Palankar, I. Panse, P. Peterson, C. Peth, J. Petriz, D. Philips, W. Pickl, S. Piconese, M. Pinti, A. G. Pockley, M. J. Podolska, C. Pucillo, S. A. Quataert, T. R. D. J. Radstake, B. Rajwa, J. A. Rebhahn, D. Recktenwald, E. B. M. Remmerswaal, K. Rezvani, L. G. Rico, J. P. Robinson, C. Romagnani, A. Rubartelli, B. Ruckert, J. Ruland, S. Sakaguchi, F. Sala-de-Oyanguren, Y. Samstag, S. Sanderson, B. Sawitzki, A. Scheffold, M. Schiemann, F. Schildberg, E. Schimisky, S. A. Schmid, S. Schmitt, K. Schober, T. Schüler, A. R. Schulz, T. Schumacher, C. Scotta, T. V. Shankey, A. Shemer, A. K. Simon, J. Spidlen, A. M. Stall, R. Stark, C. Stehle, M. Stein, T. Steinmetz, H. Stockinger, Y. Takahama, A. Tarnok, Z. Tian, G. Toldi, J. Tornack, E. Traggiai, J. Trotter, H. Ulrich, M. van der Braber, R. A. W. van Lier, M. Veldhoen, S. Vento-Asturias, P. Vieira, D. Voehringer, H. D. Volk, K. von Volkmann, A. Waisman, R. Walker, M. D. Ward, K. Warnatz, S. Warth, J. V. Watson, C. Watzl, L. Wegener, A. Wiedemann, J. Wienands, G. Willimsky, J. Wing, P. Wurst, L. Yu, A. Yue, Q. Zhang, Y. Zhao, S. Ziegler, and J. Zimmermann. 2017. Guidelines for the use of flow cytometry and cell sorting in immunological studies. Eur J Immunol 47: 1584–1797. doi: 10.1002/eji.201646632

17. Ahmed, R., L. Roger, P. Costa Del Amo, K. L. Miners, R. E. Jones, L. Boelen, T. Fali, M. Elemans, Y. Zhang, V. Appay, D. M. Baird, B. Asquith, D. A. Price, D. C. Macallan, and K. Ladell. 2016. Human Stem Cell-like Memory T Cells Are Maintained in a State of Dynamic Flux. Cell Rep 17: 2811–2818. doi: 10.1016/j.celrep.2016.11.037

18. Yu, W., N. Jiang, P. J. Ebert, B. A. Kidd, S. Müller, P. J. Lund, J. Juang, K. Adachi, T. Tse, M. E. Birnbaum, E. W. Newell, D. M. Wilson, G. M. Grotenbreg, S. Valitutti, S. R. Quake, and M. M. Davis. 2015. Clonal Deletion Prunes but Does Not Eliminate Self-Specific αβ CD8(+) T Lymphocytes. Immunity 42: 929–941. doi: 10.1016/j.immuni.2015.05.001

19. Gordon, C. L., M. Miron, J. J. Thome, N. Matsuoka, J. Weiner, M. A. Rak, S. Igarashi, T. Granot, H. Lerner, F. Goodrum, and D. L. Farber. 2017. Tissue reservoirs of antiviral T cell immunity in persistent human CMV infection. J Exp Med 214: 651–667. doi: 10.1084/jem.20160758

20. Aslan, N., L. B. Watkin, A. Gil, R. Mishra, F. G. Clark, R. M. Welsh, D. Ghersi, K. Luzuriaga, and L. K. Selin. 2017. Severity of Acute Infectious Mononucleosis Correlates with Cross-Reactive Influenza CD8 T-Cell Receptor Repertoires. MBio 8: 10.1128/mBio.01841-17

21. Sprent, J., and J. F. Miller. 1972. Interaction of thymus lymphocytes with histoincompatible cells. II. Recirculating lymphocytes derived from antigen-activated thymus cells. Cell Immunol 3: 385–404. doi:

22. Siracusa, F., M. A. McGrath, P. Maschmeyer, M. Bardua, K. Lehmann, G. Heinz, P. Durek, F. F. Heinrich, M. F. Mashreghi, H. D. Chang, K. Tokoyoda, and A. Radbruch. 2018. Nonfollicular reactivation of bone marrow resident memory CD4 T cells in immune clusters of the bone marrow. Proc Natl Acad Sci U S A 10.1073/pnas.1715618115

23. Jones, G. W., and S. A. Jones. 2016. Ectopic lymphoid follicles: inducible centres for generating antigen-specific immune responses within tissues. Immunology 147: 141–151. doi: 10.1111/imm.12554

24. Kastenmüller, W., M. Brandes, Z. Wang, J. Herz, J. G. Egen, and R. N. Germain. 2013. Peripheral prepositioning and local CXCL9 chemokine-mediated guidance orchestrate rapid memory CD8+ T cell responses in the lymph node. Immunity 38: 502–513. doi: 10.1016/j.immuni.2012.11.012

25. Darzynkiewicz, Z., L. Staiano-Coico, and M. R. Melamed. 1981. Increased mitochondrial uptake of rhodamine 123 during lymphocyte stimulation. Proc Natl Acad Sci U S A 78: 2383–2387. doi:

26. Nusse, M., W. Beisker, C. Hoffmann, and A. Tarnok. 1990. Flow cytometric analysis of G1- and G2/M-phase subpopulations in mammalian cell nuclei using side scatter and DNA content measurements. Cytometry 11: 813–821. doi: 10.1002/cyto.990110707

27. Okhrimenko, A., J. R. Grun, K. Westendorf, Z. Fang, S. Reinke, P. von Roth, G. Wassilew, A. A. Kuhl, R. Kudernatsch, S. Demski, C. Scheibenbogen, K. Tokoyoda, M. A. McGrath, M. J. Raftery, G. Schonrich, A. Serra, H. D. Chang, A. Radbruch, and J. Dong. 2014. Human memory T cells from the bone marrow are resting and maintain long-lasting systemic memory. Proc Natl Acad Sci U S A 111: 9229–9234. doi: 10.1073/pnas.1318731111

28. Kamphorst, A. O., R. N. Pillai, S. Yang, T. H. Nasti, R. S. Akondy, A. Wieland, G. L. Sica, K. Yu, L. Koenig, N. T. Patel, M. Behera, H. Wu, M. McCausland, Z. Chen, C. Zhang, F. R. Khuri, T. K. Owonikoko, R. Ahmed, and S. S. Ramalingam. 2017. Proliferation of PD-1+ CD8 T cells in peripheral blood after PD-1-targeted therapy in lung cancer patients. Proc Natl Acad Sci U S A 114: 4993–4998. doi: 10.1073/pnas.1705327114

29. Di Rosa, F. 2016. Maintenance of memory T cells in the bone marrow: survival or homeostatic proliferation? Nat Rev Immunol 16: 271. doi: 10.1038/nri.2016.31

30. Sercan Alp, O., and A. Radbruch. 2016. The lifestyle of memory CD8(+) T cells. Nat Rev Immunol 16: 271. doi: 10.1038/nri.2016.32

31. Colloca, S., E. Barnes, A. Folgori, V. Ammendola, S. Capone, A. Cirillo, L. Siani, M. Naddeo, F. Grazioli, M. L. Esposito, M. Ambrosio, A. Sparacino, M. Bartiromo, A. Meola, K. Smith, A. Kurioka, G. A. O’Hara, K. J. Ewer, N. Anagnostou, C. Bliss, A. V. Hill, C. Traboni, P. Klenerman, R. Cortese, and A. Nicosia. 2012. Vaccine vectors derived from a large collection of simian adenoviruses induce potent cellular immunity across multiple species. Sci Transl Med 4: 115ra2. doi: 10.1126/scitranslmed.3002925

32. Quinci, A. C., S. Vitale, E. Parretta, A. Soriani, M. L. Iannitto, M. Cippitelli, C. Fionda, S. Bulfone-Paus, A. Santoni, and F. Di Rosa. 2012. IL-15 inhibits IL-7Ralpha expression by memory-phenotype CD8(+) T cells in the bone marrow. Eur J Immunol 42: 1129–1139. doi: 10.1002/eji.201142019

33. Villarroya-Beltri, C., C. Gutiérrez-Vázquez, F. Sánchez-Madrid, and M. Mittelbrunn. 2013. Analysis of microRNA and protein transfer by exosomes during an immune synapse. Methods Mol Biol 1024: 41–51. doi: 10.1007/978-1-62703-453-1_4

34. Deng, N., J. M. Weaver, and T. R. Mosmann. 2014. Cytokine diversity in the Th1-dominated human anti-influenza response caused by variable cytokine expression by Th1 cells, and a minor population of uncommitted IL-2+IFNγ- Thpp cells. PLoS One 9: e95986. doi: 10.1371/journal.pone.0095986

